# Endogenous attention is invariant to sequential effects in performance around the visual field

**DOI:** 10.64898/2026.07.18.739376

**Authors:** Hsing-Hao Lee, Marisa Carrasco

## Abstract

Visual perception is shaped by both current sensory inputs and contextual information across space and time, including the perception of preceding trials. Covert spatial attention selects and prioritizes relevant sensory information. Endogenous attention is voluntary, goal-driven, and flexible. However, it cannot alleviate visual polar-angle asymmetries, specifically, the horizontal-vertical anisotropy and the vertical meridian asymmetry. Although previous studies reported sequential effects whereby attention interacts with response repetitions, it is unknown whether and how endogenous attentional modulation of performance varies as a function of target location and trial history. Here, we reanalyzed data from three published studies of endogenous attention in orientation discrimination tasks, to (1) assess typical sequential effects on response, in which response to the current trial is biased toward the previous one, and (2) examine whether sequential effects modulate performance across visual field locations, along four dimensions: (1) target location, (2) target feature, (3) attention condition, and (4) the correctness of the preceding (*n –* 1) trial. First, we observed typical sequential response repetition effects tied to the repeated location and feature properties of the target. Second, whereas attention robustly modulated performance, none of the three studies showed evidence of sequential attention effects on performance across any of the four dimensions, nor did trial history reveal any attention-by-location interactions. Together, these findings provide compelling evidence that visual polar-angle asymmetries are resistant to endogenous attention, and that even sequential effects–involving bottom up and top-down processes–do not mitigate these performance asymmetries.

## Introduction

Visual perception depends on sensory input and is modulated by its location across the visual field, as well as by sequential history and covert attention. In this study, we investigated whether and how sequential effects influence endogenous attentional modulation across the visual field, particularly around polar angle.

Visual perception depends not only on current sensory input but also on contextual information over space and time (Cicchini et al., 2024; Fecteau & Munoz, 2003; Fischer & Whitney, 2014; Kiyonaga et al., 2017; Manassi et al., 2023; Manassi & Whitney, 2024; Pascucci et al., 2023). To combat environmental noise and stabilize visual input, perception and responses at the present moment are biased toward input from the recent past. For example, the perception of an oriented grating in the current trial is biased toward the orientation of the grating in the previous trial. This phenomenon—serial dependence or the sequential effect—has been shown in orientation (Bliss et al., 2023; Can & Collins, 2025; Ceylan et al., 2021; Cicchini et al., 2021; Collins, 2019; Fischer & Whitney, 2014; Fritsche et al., 2017; Gallagher & Benton, 2022; Galluzzi et al., 2022; Gekas et al., 2019; Kondo et al., 2022; Lau & Maus, 2019; Liberman et al., 2016; Little & Clifford, 2025; Murai & Whitney, 2021; Pascucci et al., 2019; Samaha et al., 2019; Shan & Postle, 2022; St. John-Saaltink et al., 2016; Teng et al., 2022; Van Bergen & Jehee, 2019), color (Bays et al., 2009; Collins, 2022; Guan & Goettker, 2024; Maljkovic & Nakayama, 1994; Manassi et al., 2019; Yashar et al., 2013), motion (Czoschke et al., 2019; Fischer et al., 2020; Moon et al., 2023), face perception (Hsu & Lee, 2016; Kondo et al., 2022; Liberman et al., 2014; Liberman et al., 2018; Turbett et al., 2022; Van der Burg et al., 2021; Yu & Ying, 2021), and attractiveness evaluation (Ho & Newell, 2020; Kim et al., 2019; Pegors et al., 2015; Xia et al., 2016) tasks, as well as in gaze direction (Alais et al., 2018; Bliss et al., 2023; Goettker & Stewart, 2024). Some studies refer to “sequential effects”, whereas others refer to “serial dependence”. Because the former term encompasses the latter, we use ‘sequential effects’ throughout this paper.

Visual perception is also modulated by covert attention. To process rich and complex visual environments effectively, covert attention allocates limited bioenergetic resources according to task demands in the absence of gaze shifts (Carrasco, 2011; Lennie, 2003; Pestilli & Carrasco, 2005). Endogenous attention is voluntary, goal-driven, flexible, and sustained, whereas exogenous attention is involuntary, stimulus-driven, automatic, and transient. Both improve performance (for reviews, Carrasco, 2011; Carrasco & Barbot, 2014) and alter appearance (for a review, Carrasco & Barbot, 2019) across a wide range of visual tasks. Endogenous attention is more flexible than exogenous attention, modulating perceptual sensitivity adaptively as a function of task demands. For example, it enhances contrast sensitivity (Jigo & Carrasco, 2020) and increases gain (Fernández et al., 2022) across a wider range of spatial frequencies than exogenous attention. Furthermore, endogenous attention improves performance in texture segregation tasks regardless of spatial resolution (Barbot & Carrasco, 2017; Yeshurun et al., 2008), whereas exogenous attention can improve or impair performance depending on spatial resolution (Carrasco et al., 2006; Yeshurun & Carrasco, 1998). It also facilitates temporal-order judgments, whereas exogenous attention can impair them (Hein et al., 2006). Finally, endogenous attention scales with cue validity to allocate resources accordingly, but exogenous attention does not (Giordano et al., 2009). Thus, we investigated whether and how the flexibility of endogenous attention extends to trial-by-trial adjustments of sequential effects and varies across the visual field.

Visual performance is not homogeneous around the visual field; in human adults, it is better at the horizontal than the vertical meridian (horizontal-vertical anisotropy, HVA) and better at the lower than the upper vertical meridian (vertical meridian asymmetry, VMA). These performance fields are present in many tasks, including contrast sensitivity (Abrams et al., 2012; Baldwin et al., 2012; Cameron et al., 2002; Carrasco et al., 2001; Corbett & Carrasco, 2011; Himmelberg et al., 2020; Kwak, Lu, & Carrasco, 2024; Lee & Carrasco, 2025), visual acuity (Barbot & Carrasco, 2017; Kwak et al., 2023), spatial resolution (Altpeter et al., 2000; Carrasco et al., 2002; Greenwood et al., 2017), and motion (Fuller & Carrasco, 2009; Tünçok, Kiorpes, & Carrasco, 2025). They also emerge in mid-level processes such as texture segregation (Talgar & Carrasco, 2002; Wang et al., 2020), size perception (Schwarzkopf, 2019), and crowding (Greenwood et al., 2017; Kurzawski et al., 2023; Petrov & Meleshkevich, 2011), as well as high-level tasks such as letter recognition (Mackeben, 1999), numerosity perception (Chakravarthi et al., 2022), face perception (Afraz et al., 2010; Kim & Chong, 2024), word identification (Tsai et al., 2024), and visual short-term memory (Montaser-Kouhsari & Carrasco, 2009).

Performance fields are not easily reshaped; despite its flexibility, endogenous attention benefits performance at all locations to a similar degree and does not compensate for these asymmetries (Lee & Carrasco, 2026; Purokayastha et al., 2021; Tünçok, Carrasco, & Winawer, 2025). Performance fields are also resistant to exogenous attention (Cameron et al., 2002; Roberts et al., 2016, 2018) and endogenous temporal attention (Fernández et al., 2019). Moreover, presaccadic attention, which enhances processing at the impending saccade target, can even exacerbate them by enhancing contrast sensitivity most at the horizontal meridian and least at the upper vertical meridian (Hanning et al., 2022, 2024; Kwak et al., 2025; Kwak, Zhao, et al., 2024).

The relation between covert spatial attention and sequential effects remains unclear. Some studies reported that sequential effects decrease when no attention is allocated to previously experienced stimuli (Fischer & Whitney, 2014; Fritsche & de Lange, 2019; Rafiei, Hansmann-Roth, et al., 2021), whereas others reported an absence of such dependence (Collins, 2022; Fornaciai & Park, 2018; Goettker & Stewart, 2022). However, these studies did not monitor eye movements or cleanly isolate endogenous from exogenous attention (Carrasco, 2011;

Carrasco & Barbot, 2014; Chica et al., 2013; Olivers, 2025), rendering their findings inconclusive. The few studies that have explored sequential effects with properly operationalized attention have shown similar sequential effects in task performance, but differential oculomotor responses. For example, studies on presaccadic attention showed that performance in orientation discrimination tasks remains stable regardless of whether participants performed spontaneous or cue-driven saccades on the previous trial (Tyralla & Zimmermann, 2025). However, performance improves when the saccade on the previous trial was directed to the same location (White et al., 2013). Likewise, when deploying covert attention to a specific time point, the effects of temporal attention on performance are similar across trials regardless of temporal attention and expectations on the previous trial (Duyar & Carrasco, 2025b). However, microsaccades are biased by expectations formed on the previous trial (Duyar & Carrasco, 2025a).

Sequential effects may be stronger at peripheral than central locations (Kandemir & Olivers, 2026, but see Suárez-Pinilla et al., 2018). Given that perceptual sensitivity decreases with eccentricity (for reviews, Himmelberg et al., 2023; Olivers, 2025; Strasburger et al., 2011), this peripheral increase suggests a heavier reliance on prior experience when perceptual sensitivity decreases. Whether this explanation generalizes to polar angle differences remains unknown, as different mechanisms underlie sensitivity variations across eccentricity and around polar angle (Jigo et al., 2023; Xue et al., 2024; Xue, Barbot, et al., 2026; Xue, Landy, & Carrasco, 2026).

To investigate whether and how sequential effects influence endogenous attentional modulation around polar angle, we assessed sequential effects in three published studies of orientation discrimination in terms of (1) response repetition and (2) perceptual sensitivity (**Figure 1**). Although stimulus parameters varied across these studies (e.g., stimulus size, eccentricity, orientations, and number of possible target locations; see Methods for details), we evaluated sequential effects along three key dimensions (Manassi et al., 2023; Manassi & Whitney, 2024): (1) spatial: as spatial distance between successive targets increases, sequential effects decrease (Collins, 2019; Fischer & Whitney, 2014; Fornaciai & Park, 2018; Luo et al., 2022); (2) featural: as feature similarity between consecutive targets decreases, sequential effects decrease (Barbosa & Compte, 2020; Fischer & Whitney, 2014; Liberman et al., 2014; Turbett et al., 2021); and (3) attentional, with studies reporting stronger sequential effects produced by valid than invalid attention conditions in the previous target (Fischer & Whitney, 2014; Fornaciai & Park, 2018; Fritsche & de Lange, 2019; Kim et al., 2020; Liberman et al., 2016; Rafiei, Chetverikov, et al., 2021; Rafiei, Hansmann-Roth, et al., 2021).

**Figure 1.**
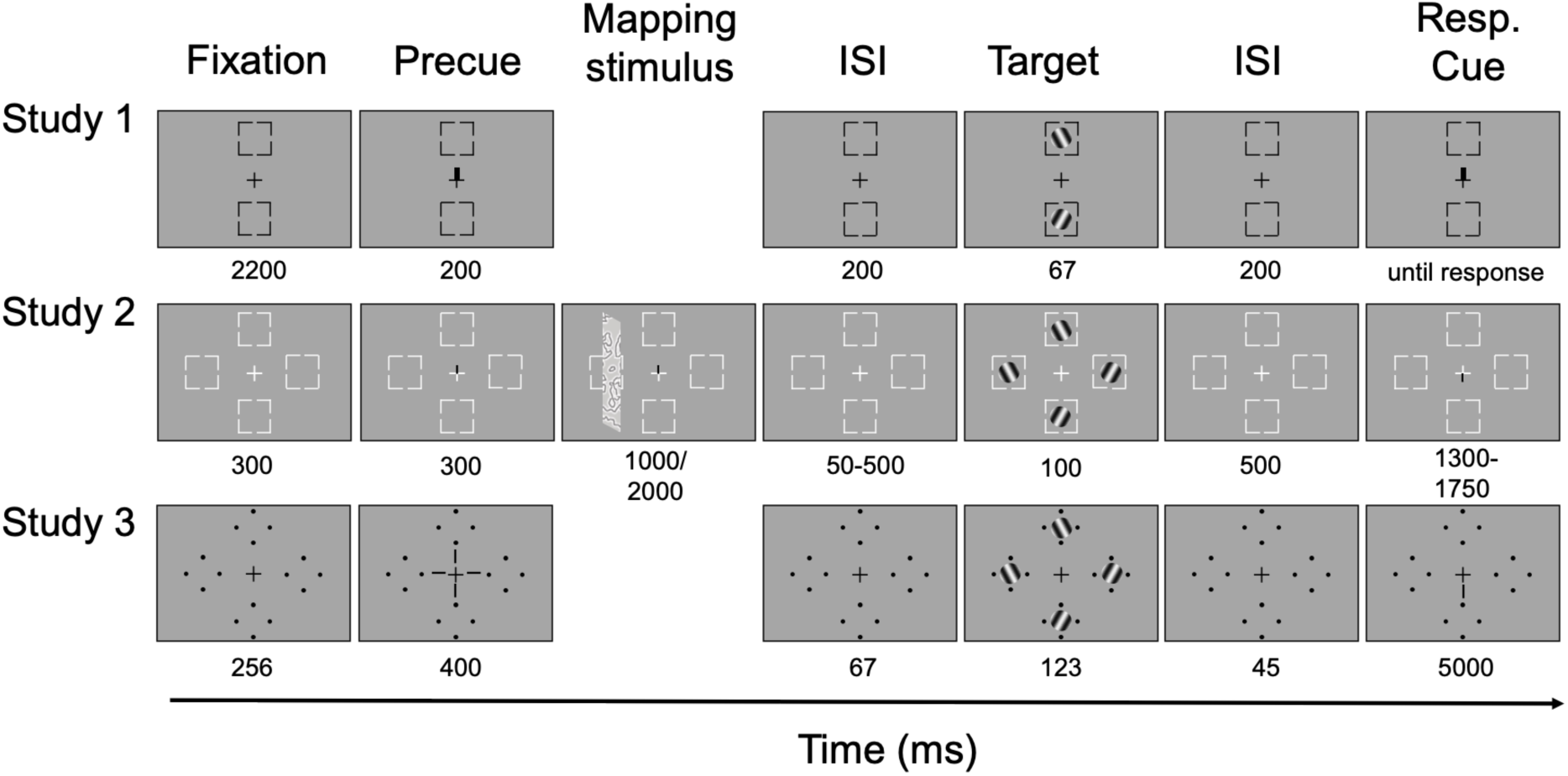
Trial sequence: Panels illustrate example trials for valid (Study 1, top panel), invalid (Study 2, middle panel), and neutral (Study 3, bottom panel) cue conditions, determined by the relation between the precue and the response cue location. Numbers below each frame indicate duration (in ms). Stimulus size, spatial frequency, and background luminance are not to scale.

Furthermore, we examined whether sequential effects depend on the correctness of the previous (*n –*1) trial. Given the flexible nature of endogenous attention, participants may adapt strategies based on prior performance. On the one hand, they may deploy more resources to the target when the previous trial was incorrect than when it was correct (Danielmeier & Ullsperger, 2011; Little & Clifford, 2025). This compensatory mechanism may enhance performance more at locations with poorer sensitivity (i.e., the upper vertical meridian). On the other hand, post-error improvement in accuracy does not always occur (Adkins et al., 2024; Danielmeier & Ullsperger, 2011; Ullsperger et al., 2014).

The aims of this study were twofold: First, we assessed typical sequential effects in response repetition (i.e., response bias), in which the response to the current trial is biased toward the previous (*n –*1) trial. Second, and more importantly, we examined whether sequential effects modulate performance across visual field locations in endogenous attention tasks.

## EXPERIMENTS

### Materials and Methods

This study reanalyzed data from three published studies. All participants had normal or corrected-to-normal vision. The Institutional Review Board at New York University approved the experimental procedures, and all participants provided informed consent before participating. We analyzed data from 12 adults (mean age: 28 years), including author HHL, who participated in Study 1 (Lee & Carrasco, 2026); nine adults (mean age: 28.5 years) who participated in Study 2 (Tünçok, Carrasco, & Winawer, 2025); and 18 of the 20 participants (mean age: 27.3 years, data from the remaining two participants could not be recovered) in Experiment 2 of Study 3 (Purokayastha et al., 2021).

### Stimuli and Apparatus

Stimuli in all experiments were generated using MATLAB (MathWorks, Natick, MA) and the Psychophysics Toolbox (Psychtoolbox-3; Brainard, 1997; Pelli, 1997), and eye movements were monitored using an EyeLink 1000 eye tracker (SR Research, Ottawa, ON, Canada) at a sampling rate of 1,000 Hz to ensure fixation. Other details are specified below as well as in the original publications.

#### Study 1

The target Gabor (4° diameter, 5 cycles per degree (cpd), 1.25° full width at half maximum) was presented at four cardinal locations (left, right, upper, and lower), and the distractor Gabor was presented at the opposite side of the meridian (see below), each positioned 8° from fixation (center-to-center). Four black placeholders (0.16° length × 0.06° width) were located 0.5° from the edge of the Gabor. A central fixation cross, consisting of a plus sign (0.25° length × 0.06° width), remained on the screen throughout the trial. The endogenous attentional cue (0.75° length × 0.2° width) was presented at fixation. Each participant completed two sessions.

Participants were seated in a dimly lit, sound-attenuated room, with their heads stabilized with a chinrest at a viewing distance of 57 cm. Stimuli were displayed on a gamma-corrected 20-inch ViewSonic G220fb CRT monitor (ViewSonic Corporation, Brea, CA) with a resolution of 1280 × 960 pixels and a refresh rate of 100 Hz.

#### Study 2

The target Gabor (3° diameter, 4 cpd, 1° full width at half maximum) was presented at four cardinal locations (left, right, upper, and lower), each positioned 6° from fixation (center-to-center). Four white placeholders (0.22° length × 0.05° width) were located 0.11° from the edge of the Gabor. A white central fixation cross, consisting of a plus sign (0.35° length × 0.02° width), remained on the screen throughout the trial. The endogenous attentional cue was black and occupied one of the four arms of the fixation cross.

Participants lay supine in a 3-T Siemens MAGNETOM Prisma MRI scanner (Siemens Medical Solutions, Erlangen, Germany); their heads were secured with pads to reduce head motion. Participants viewed the screen through an angled mirror mounted on the head coil. The total viewing distance (eye to mirror plus mirror to screen) was 86.5 cm. Stimuli were displayed on a ProPixx DLP LED projector with a linear gamma table (VPixx Technologies Inc., Saint-Bruno-de-Montarville, QC, Canada). The projected screen (60 × 36.2 cm) had a resolution of 1920 × 1080 pixels with a refresh rate of 60 Hz.

#### Study 3

The target Gabor (3.2° diameter, 4 cpd, ∼1° full width at half maximum) was presented at four cardinal locations (left, right, upper, and lower), each positioned 6.4° from fixation (center-to-center). Four placeholders, each composed of four black dots (0.05° radius), were arranged in a circle 0.5° from the edge of the upcoming Gabor patch stimulus. A central fixation cross, consisting of a plus sign (0.5° length), remained on the screen throughout the trial. The endogenous attentional cue was either a single 0.88° line or four 0.28° lines (all 0.14° thick), which were positioned 0.38° from the center of the fixation cross to indicate the possible target locations.

Participants were seated in a dimly lit, sound-attenuated room, with their heads stabilized with a chinrest at a viewing distance of 57 cm. Stimuli were displayed on a gamma-corrected 21-inch IBM P260 CRT monitor with a resolution of 1280 × 960 pixels and a refresh rate of 90 Hz.

### Psychophysical Tasks and Procedures

All three tasks followed the same endogenous attention protocol but with different timing parameters (as specified in **Figure 1**). We titrated stimulus contrast (Studies 1 and 3) or tilt angle (Study 2) to equate stimulus discriminability across locations.

#### Study 1

In the endogenous attention task, each trial began with 2,200 ms of fixation, followed by an endogenous precue presented for 100 ms and a 200-ms inter-stimulus interval (ISI). Two tilted Gabor patches were then presented for 67 ms, followed by a second 200-ms ISI and a response cue. In valid trials, the location indicated by the response cue matched that of the precue. In invalid trials, the locations mismatched. In neutral trials the precue pointed to both locations. Participants judged whether the target Gabor was tilted clockwise or counterclockwise from vertical (±2.5°). Participants were instructed to respond as accurately as possible while maintaining central fixation throughout the trial. A trial would be interrupted and repeated at the end of the block if a participant’s eye position deviated ≥1.5° from the center at any point between precue onset until response-cue onset. Feedback regarding accuracy was provided after each trial.

Overall, 20% of the trials contained a neutral cue, whereas the remaining 80% contained an informative attentional cue pointing toward a specific location (75% valid, 25% invalid). Gabor patch locations were blocked in horizontal-meridian and vertical-meridian sessions with the order of these two sessions randomized across participants. There were 400 trials per location, 1,600 trials in total. Stimulus orientation was fully counterbalanced, yielding an equal number of clockwise (CW: 50%) and counterclockwise (CCW: 50%) trials.

To equate stimulus discriminability across locations, before the endogenous attention task, participants’ contrast thresholds were titrated. There were four independent staircases per location, varying Gabor contrast from 2% to 85% to reach ∼75% accuracy in the orientation discrimination task. Each staircase started at different contrast levels (85%, 2%, the median contrast of 43.5%, or a random value between 2% and 85%) and contained 48 trials. Four blocks (192 trials per location) were conducted consecutively for either the horizontal meridian or the vertical meridian. Contrast thresholds were derived using an adaptive staircase procedure implemented in the Palamedes toolbox (Prins & Kingdom, 2018). The Gabors in the thresholding procedure were always preceded by a neutral precue, which provided the same temporal information as the valid and invalid cues without providing spatial information (as was also the case in Studies 2 and 3).

#### Study 2

In the endogenous attention task, trials began with 300 ms of central fixation, after which the fixation cross changed into a precue for 300 ms followed by a mapping stimulus presented for 1,000 or 2,000 ms with the attentional precue remaining on the screen. The mapping stimulus was followed by a jittered ISI of 50 ms (10% of trials) or 300, 400, or 500 ms (30% of trials each), with only the white fixation cross presented on the screen. Then, four tilted Gabor stimuli (10% contrast) were presented for 100 ms, followed by a 500-ms ISI and a response cue. Participants judged whether the target Gabor was tilted clockwise or counterclockwise from horizontal. Participants had a 1,350*–*1,700 ms response window (to accommodate trial synchronization with MRI volume acquisition). Participants were asked to respond as accurately as possible while fixating at the center of the screen throughout the trial. Accuracy feedback was provided after each trial.

Neutral cues –pointing to all possible target locations– were presented on 20% of trials; the remaining 80% contained an informative attentional cue pointing to a specific location (75% valid, 25% invalid). Seven participants completed 2,080 trials (across four scan sessions), and two participants completed 1,560 trials (across three scan sessions). Target Gabor orientation (CW, CCW) was randomly assigned; there were 52.76% CW trials and 47.24% CCW trials on average.

To equate stimulus discriminability across target locations, before the endogenous attention task, participants’ tilt angle thresholds were titrated. Thresholds were defined as the tilt deviation corresponding to 82% accuracy on a fitted Weibull function estimated using a Best PEST adaptive procedure with 3-down/1-up staircases interleaved at each target location. The thresholding comprised about 65 trials at each target location (∼260 trials). The thresholding session also provided feedback informing the participants if their gaze deviated from the fixation cross (≥2°) by a change in the fixation cross color from white to blue.

#### Study 3

In the endogenous attention task, trials began with a 256-ms central fixation, an endogenous precue was presented for 400 ms followed by a 67-ms ISI. Four tilted Gabors were then presented for 123 ms, followed by another 45-ms ISI and a 667-ms response cue. Participants judged whether the target Gabor was tilted clockwise or counterclockwise from vertical (±20°). Participants were instructed to respond as accurately as possible while maintaining central fixation throughout the trial. A trial was interrupted and repeated at the end of the block if a participant’s eye position deviated ≥1° from the center between trial initiation and stimulus offset. Feedback regarding accuracy was provided after each trial.

The endogenous attention task contained 80 trials per block across approximately 12 blocks yielding approximately 120 trials per location in each attention condition. The target Gabor orientation (CW or CCW) was randomly assigned (49.71% CW trials and 50.29% CCW trials).

To equate stimulus discriminability across locations, before the endogenous attention task, participants’ contrast thresholds were titrated. A PEST staircase procedure involving eight interleaved 3-down/1-up staircases (two per location; one starting at 75% and the other at 25%) was used to estimate individual contrast thresholds (∼80% overall accuracy).

### Statistical Analysis

We quantified the probability of a participant giving a repeated response based on whether the previous (*n* – 1) trial was congruent (repeated) or incongruent (different) with the current trial. Task performance was indexed by *d′* [*z*(hit rate) – *z*(false alarm rate)] across conditions. Correct discrimination of clockwise trials was considered “hits,” and incorrect discrimination of counterclockwise trials was considered “false alarms,” consistent with previous studies (e.g., Jigo & Carrasco, 2020; Lee et al., 2024; Zhang et al., 2019). Bayes factors were estimated using the *BayesFactor* package in R (Morey et al., 2015) for 3-way interactions to evaluate evidence for or against an interaction among attention, polar angle location, and the sequential factor. Default Jeffreys–Zellner–Siow (JZS) priors were used, with fixed effects assigned a medium scale (*r*=0.5) and random effects a nuisance scale (*r*=1). *BF_10_* values smaller than 0.33, 0.1, and 0.01 indicate moderate, strong, and very strong evidence favoring the null hypothesis, respectively (Raftery, 1999).

To control the family-wise error rate across the four sequential-effect analyses, we applied the Holm correction (Holm, 1979) separately within each study. Specifically, for each ANOVA effect, we jointly corrected the four *p*-values obtained from analyses based on previous-trial location, orientation, attentional condition, and response correctness. For example, the four *p*-values testing the main effect of trial history were corrected as one family, as were those testing the history × location interaction and the history × current attention × location interaction.

## Results

### Sequential Effects on Response Repetition

#### Study 1

In this experiment (Lee & Carrasco, 2026), target locations were blocked along either the vertical or the horizontal meridian; thus, each trial had two possible target locations. Participants also performed blocks following adaptation on different days, but the adaptation sessions were not analyzed in the current study.

We excluded the first trial of each block and labeled the remaining trials based on whether the target location was the same as or different from that on the previous (*n –*1) trial. We conducted a two-way repeated-measures analysis of variance (rmANOVA) on the factors of attention condition (valid, neutral, invalid) and location repetition (same, different) while pooling all target locations in the analysis. For the location analysis (**Figure 2A, left panel**), there was a main effect of location repetition [*F*(1,11) = 16.88, *p* = .002, *p-Holm* = .005], no main effect of attention [*p* > .1], and no two-way interaction after correction for multiple comparisons [*F*(2,22) = 3.87, *p* = .036, *p-Holm* = .109].

**Figure 2.**
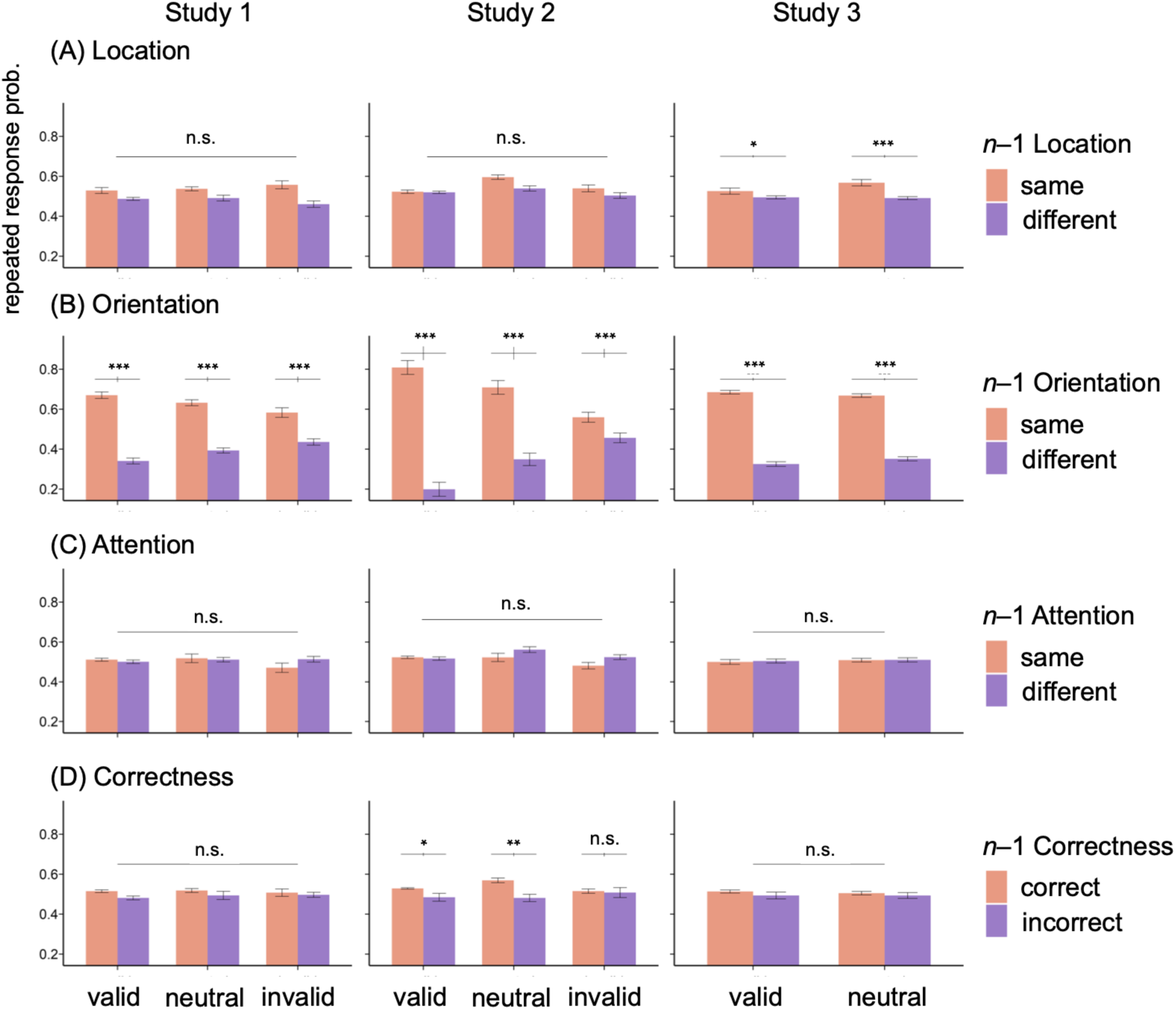
Probability of giving a repeated response in the orientation discrimination task when the (A) location, (B) target’s orientation, or (C) attentional cueing condition in the previous (*n –*1) trial was the same or different from the current trial. (D) Probability of giving the same response in the task when performance on the previous (*n –*1) trial was correct or incorrect. Error bars within the bar plots depict ±1 *SEM* of the condition; error bars above the bars indicate ±1 *SEM* of the difference between conditions.

For the feature analysis (**Figure 2B, left panel**), we conducted a two-way rmANOVA on the factors of attention (valid, neutral, invalid) and orientation repetition (same, different). There was a main effect of orientation repetition [*F*(1,11) = 104.4, *p* < .001, *p-Holm* < .001], no main effect of attention [*p*> .1] and a two-way interaction [*F*(2,22) = 31.47, *p* < .001, *p-Holm* < .001]. Paired *t*-tests showed that participants gave more repeated responses in valid [*t*(11) = 11.58, *p*<.001], neutral [*t*(11,) = 10.68, *p* < .001], and invalid trials [*t*(11) = 5.05, *p* < .001] when the previous trial was at the same orientation than the different orientation from the current one. Here, a larger probability difference between the same and different orientation trials was observed in the valid than in the neutral condition [*t*(11) = 5.36, *p* <.001], which in turn was larger in the neutral than in the invalid condition [*t*(11) = 3.43, *p* = .006].

For the attention analysis (**Figure 2C, left panel**), a two-way rmANOVA on the factors of attention (valid, neutral, invalid) and attention repetition (same, different) yielded neither main effects nor interactions [*ps* > .1]. Lastly, for the correctness analysis (**Figure 2D, left panel**), a two-way rmANOVA on the factors of attention (valid, neutral, invalid) and *n –*1 correctness (correct, incorrect) revealed neither main effects nor interactions [*ps* >.05].

#### Study 2

Study 2 was a psychophysical experiment conducted in a functional magnetic resonance imaging (fMRI) scanner (Tünçok, Carrasco, & Winawer, 2025) featuring four possible target locations. We implemented the same labeling procedure and statistical analyses as in Study 1.

We conducted a two-way rmANOVA on the factors of attention condition (valid, neutral, invalid) and location repetition (same, different) (**Figure 2A, middle panel)**. There were main effects of location repetition [*F*(1,8) = 10.56, *p* = .012, *p-Holm* = .023] and attention [*F*(2,16) = 7.94, *p* = .004, *p-Holm* = .016], but not a two-way interaction after correction for multiple comparisons [*F*(2,16) = 3.94, *p* = .041, *p-Holm* = .122].

For the feature analysis (**Figure 2B, middle panel**), we conducted a two-way rmANOVA on the factors of attention (valid, neutral, invalid) and orientation repetition (same, different). There was a main effect of orientation repetition [*F*(1,8) = 42.13, *p* < .001, *p-Holm* < .001], no main effect of attention [*p* >.1] and a two-way interaction [*F*(2,16) = 88.14, *p* < .001, *p-Holm* < .001]. Participants gave more repeated responses in valid [*t*(8) = 8.91, *p* < .001], neutral [*t*(8) = 5.7, *p* <.001], and invalid trials [*t*(8) = 2.37, *p* = .045] when the previous trial shared the same orientation than a different orientation from the current one. As in Study 1, a larger probability difference between the same and different orientation trials was observed in the valid than in the neutral condition [*t*(8) = 7.2, *p* <.001], which in turn was larger than in the invalid condition [*t*(8) = 7.92, *p* = .006].

For the attention analysis (**Figure 2C, middle panel**), we conducted a two-way rmANOVA on the factors of attention (valid, neutral, invalid) and attention repetition (same, different). Neither the main effect of attention, after correction for multiple comparisons [*F*(2,16) = 5, *p* = .021, *p-Holm* = .062], nor that of attention repetition, nor their interaction [*ps* >.05] was significant.

Lastly, for the correctness analysis (**Figure 2D, middle panel**), a two-way rmANOVA on the factors of attention (valid, neutral, invalid) and *n* –1 correctness (correct, incorrect) yielded a main effect of correctness [*F*(1,8) = 27.42, *p* < .001, *p-Holm* = .002], but no main effect of attention [*p*>.1] and no interaction after correction for multiple comparisons [*F*(2,16) = 3.52, *p* = .054, *p-Holm* = .122].

#### Study 3

Study 3 (Purokayastha et al., 2021) included only valid and neutral conditions, omitting the invalid condition. Nonetheless, we conducted the same analysis procedure as in Studies 1 and 2: a two-way rmANOVA on the factors of attention (valid, neutral) and location repetition (same, different) (**Figure 2A, right panel**). There was a main effect of location repetition [*F*(1,17) = 30.07, *p* <.001, *p-Holm* < .001], no effect of attention [*p*> .1] and a two-way interaction [*F*(1,17) = 8.12, *p* = .011, *p-Holm* = .033]. Participants gave more repeated responses in valid [*t*(17) = 2.71, *p* = .015] and neutral [*t*(17) = 5.6, *p* < .001] trials when the previous trial was at the same location rather than at a different location, which led to a larger probability difference between the same and different location conditions in the neutral than in the valid condition [*t*(17) = 2.85, *p* = .011].

For the feature analysis (**Figure 2B, right panel**), we conducted a two-way rmANOVA on the factors of attention (valid, neutral) and orientation repetition (same, different). There was a main effect of orientation repetition [*F*(1,17) = 1182, *p* < .001, *p-Holm* <.001], no main effect of attention [*p* > .1] and a two-way interaction [*F*(1,17) = 10, *p* = .006, *p-Holm* = .023]. Participants gave more repeated responses in valid [*t*(17)=26.47, *p* < .001] and neutral [*t*(17) = 31.39, *p* < .001] trials when the previous trial shared the same orientation as the current one, which led to a larger probability difference between the same and different orientation conditions in the valid than in the neutral condition [*t*(17) = 3.16, *p* = .006]—the same pattern observed in Studies 1 and 2, except that there was no invalid condition in this study.

For the attention analysis (**Figure 2C, right panel**), a two-way rmANOVA on attention (valid, neutral) and attention repetition (same, different) yielded neither main effects nor an interaction [*ps* > .1]. Lastly, for the correctness analysis (**Figure 2D, right panel**), a two-way rmANOVA on attention (valid, neutral) and *n –*1 correctness (correct, incorrect) revealed neither main effects nor an interaction [*ps* > .1].

#### Combined analysis for response repetition

To examine the consistency of our findings across studies and enhance statistical power, we combined the data from (1) Studies 1*–*3 for valid and neutral trials, and (2) Studies 1 and 2 for invalid trials, and conducted four analyses. We treated the probability of giving a repeated response as the dependent variable and conducted a series of 2-way rmANOVAs on the factors of attention (valid, neutral) and trial history— location, orientation, attention repetition (same, different), and *n* –1 correctness (correct, incorrect).

For the location repetition analysis (**Figure 3A**), results revealed main effects of location repetition [*F*(1,38) = 44.28, *p* < .001, *p-Holm* < .001] and attention [*F*(1,38) = 10.17, *p* = .003, *p-Holm* = .014], as well as an interaction [*F*(1,38) = 17.38, *p* < .001, *p-Holm* < .001]. Paired *t*-tests revealed that the probability difference in repeating a response when the previous location was the same vs. different was larger in the neutral than in the valid condition [*t*(38) = 4.16, *p* <.001]. Across Studies 1 and 2, participants gave more repeated responses when the previous location was the same than different from the current location in the invalid condition [*t*(20) = 3.96, *p* <.001].

**Figure 3.**
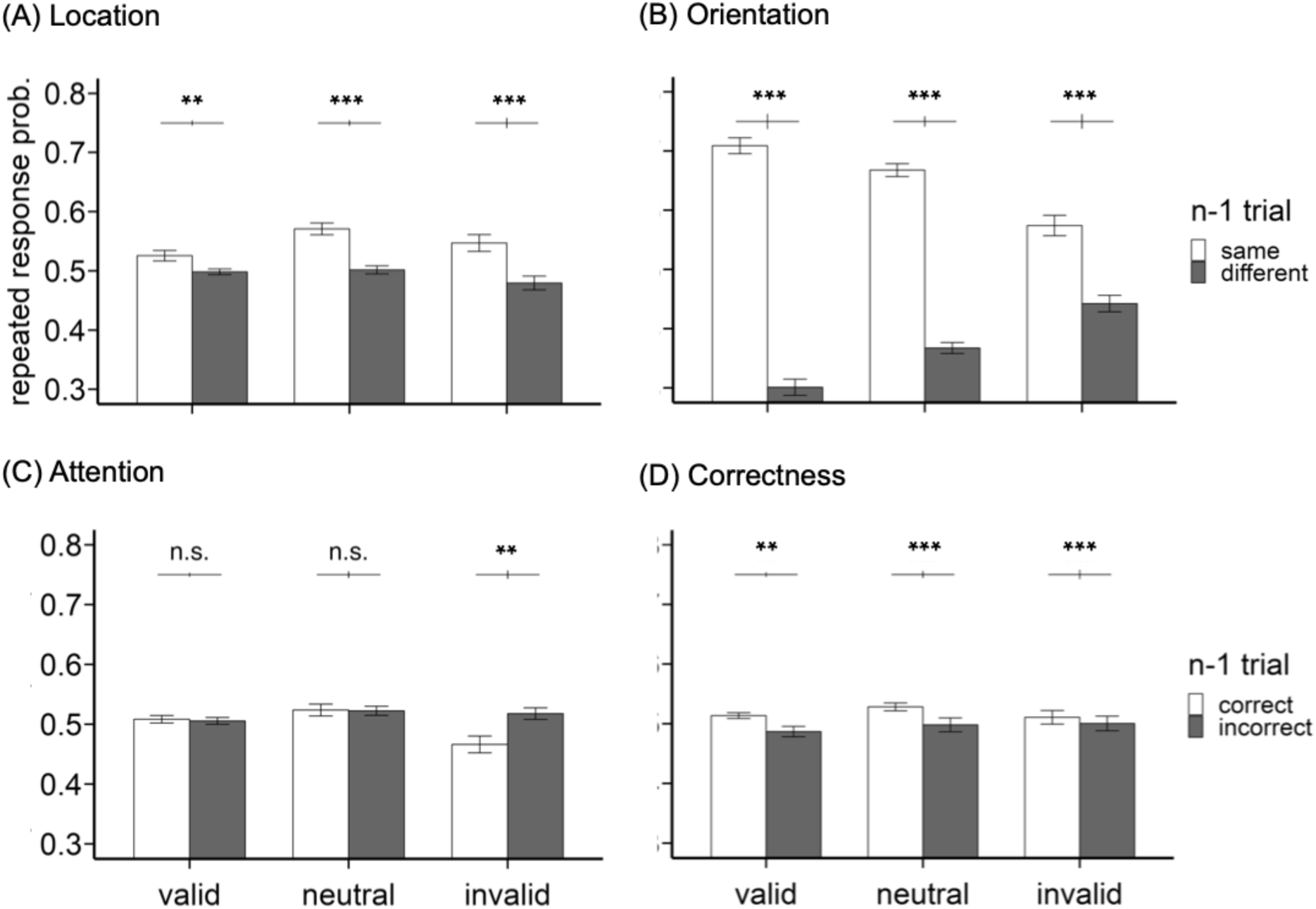
Combined analysis across Studies 1–3. Probability of giving a repeated response in the orientation discrimination task when the (A) location, (B) target’s orientation, or (C) attentional cueing condition in the previous (*n –*1) trial was the same as or different from the current trial. (D) Probability of giving the same response in the task when performance on the previous (*n –*1) trial was correct or incorrect. Error bars within the bars depict ±1 SEM of the condition; error bars above the bars indicate ±1 SEM of the difference between conditions. Valid and neutral conditions include data from Studies 1**–**3; the invalid condition includes only Studies 1 and 2, as Study 3 did not test invalid trials.

For the feature repetition analysis (**Figure 3B**), results revealed a main effect of orientation repetition [*F*(1,38) = 310.8, *p*<.001, *p-Holm* < .001], no effect of attention [*F*(1,38) = 3.83, *p* = .058, *p-Holm* = .173], and an interaction [*F*(1,38) = 37.5, *p* < .001, *p-Holm* <. 001]. Paired *t*-tests revealed that the probability difference in giving repeated responses when the previous orientation was the same vs. different was larger in the valid than in the neutral condition [*t*(38) = 6.12, *p* < .001]. Across Studies 1 and 2, participants also gave more repeated responses when the previous orientation was the same than different in the invalid condition [*t*(20) = 5.34, *p* < .001]. To ensure that these effects did not simply arise from the structure of the target feature and its corresponding responses, but rather from sequential effects, we conducted a permutation analysis in which trial order was randomly shuffled 10,000 times within each participant. For each permutation, the relation between the previous (*n* – 1) and current trial was recomputed based on the newly adjacent trials, and we calculated the difference in the probability of giving a repeated response when the trial was the same versus different orientation, D = P (response repetition ∣ same orientation) – P (response repetition ∣ different orientation). The D values were then pooled across Studies 1–3 for valid and neutral conditions, and across Studies 1–2 for the invalid condition. In the valid condition, the observed effect (D_observed_ = 0.41) was greater than that from randomly paired trials (D_shuffled_ = 0.38, permutation *p* < .001). The same result was obtained for the neutral condition (D_observed_ = 0.28, D_shuffled_ = 0.02, permutation *p* < .001), but not in the invalid condition (D_observed_ = 0.13, D_shuffled_ = 0.13, permutation *p* = .738). The findings in the valid and neutral conditions of the permutation test were in line with the combined analysis, whereas the findings in the invalid condition in the combined analysis, which could be due to the target feature itself rather than to sequential effect, is no longer significant in the permutation analysis.

For the attention repetition analysis (**Figure 3C**) across Studies 1–3, results revealed no main effect of attention after correction for multiple comparisons, [*F*(1,38) = 4.7, *p* = .036, *p-Holm* = .146], nor other significant effects [*ps* > .1]. Interestingly, in the invalid condition in Studies 1 and 2, participants gave more repeated responses when the previous trial was in a different attentional condition from the current one [*t*(20) = 3.66, *p* = .002].

We also conducted a 4-way rmANOVA for location repetition, orientation repetition, attention repetition and attention condition (valid, neutral) by pooling participants from all three studies to see if different trial history aspects interact. We observed no significant 4-way interaction [*F*(1,38) < 1], but 3-way interactions among (1) location repetition, orientation repetition, and attention condition [*F*(1,38) = 4.91, *p* = .033], and (2) orientation repetition, attention repetition, and attention condition [*F*(1,38) = 35.57, *p* < .001]. The post-hoc analyses are reported in the *Supplementary Materials*.

For the *n –*1 correctness analysis (**Figure 3D**), results revealed a main effect of correctness [*F*(1,38) = 13.18, *p* < .001, *p-Holm* = .005]; participants gave more repeated responses when the previous trial was correct than incorrect. No other effects were significant [*ps*>.1]. Because there were more trials following correct than incorrect responses given the experimental design, we conducted a down-sampling analysis in which, separately for each participant and attention condition, trials following correct responses were randomly down-sampled to match the number of trials following incorrect responses (on average, 127 trials per attention and correctness condition per participant). This procedure was repeated 1,000 times. The main effect of previous-trial correctness remained significant in 997 of the 1,000 down-sampled iterations (99.7%; median *p* = .001), indicating that the correctness effect was robust to the imbalance in trial numbers. Additionally, the main effect of attention and the correctness × attention interaction were significant in only 0.1% and 0.2% of the iterations, respectively (median *ps >* .1), supporting the combined analysis without down-sampling. There was no difference in the probability of giving a repeated response in the invalid condition for correct and incorrect trials, either without [*t*(20) = 0.66, *p* > .1] or with [median *p* > .1] down-sampling (on average, 76.9 trials per correctness condition per participant).

### Sequential Effects on Performance

#### Study 1

To analyze whether target location repetition enhanced orientation perception (**Figure 4A**), we excluded the first trial of each block and labeled the remaining trials based on whether the target location was the same as or different from that on the previous (*n –*1) trial. We conducted a 3-way rmANOVA on the factors of attention condition (valid, neutral, invalid), location (left, right, upper, lower), and location repetition (same, different). Results revealed a main effect of attention [*F*(2,22) = 51.91, *p* < .001, *p-Holm* < .001], but neither a main effect of location nor location repetition [*ps* > .1] nor any two-way [*ps* > .05] or three-way [*p* > .1, *BF_10_* = 0.05] interactions.

**Figure 4.**
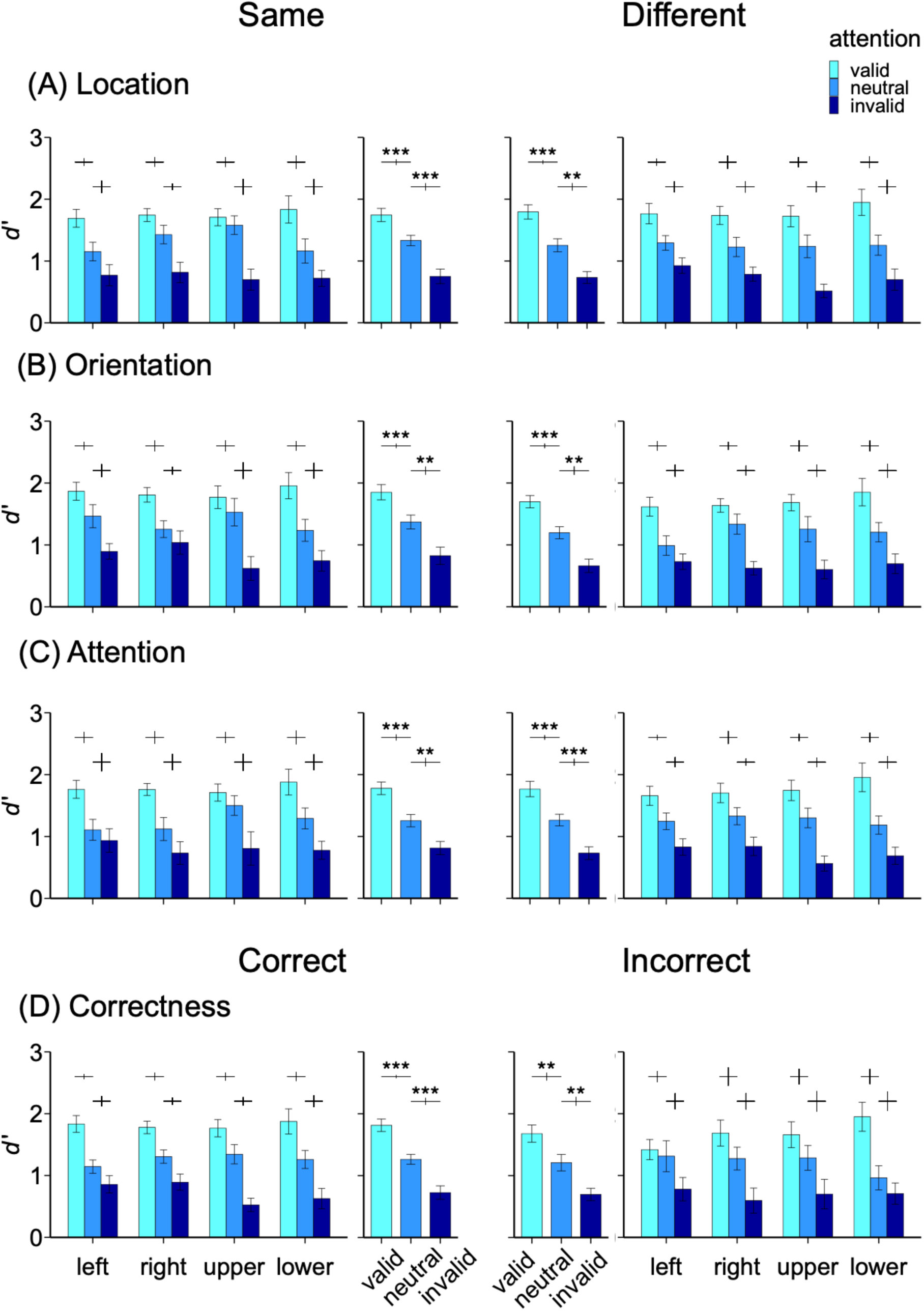
Study 1. Sensitivity in the orientation discrimination task when the (A) location, (B) target’s orientation, or (C) attentional cueing condition in the previous (*n –* 1) trial was the same as or different from the current trial. (D) Sensitivity when performance on the previous (*n –*1) trial was correct or incorrect. For each panel, left and right bars represent the same and different (or correct and incorrect) conditions, respectively. The middle two columns indicate the results averaged across locations. Error bars within the bars depict ±1 *SEM* of the condition; error bars above the bars indicate ±1 *SEM* of the difference between conditions.

For the feature analysis (**Figure 4B**), we conducted a 3-way rmANOVA on the factors of attention (valid, neutral, invalid), location (left, right, upper, lower), and orientation repetition (same, different). Results revealed a main effect of attention [*F*(2,22) = 51.17, *p* < .001, *p-Holm* < .001], but neither a main effect of location nor orientation repetition [*ps* > .05], nor any two-way [*ps* > .1] or three-way [*p* > .1, *BF_10_* = 0.13] interactions.

For the attention analysis (**Figure 4C**), we conducted a 3-way rmANOVA on the factors of attention (valid, neutral, invalid), location (left, right, upper, lower), and attention repetition (same, different). Results revealed a main effect of attention [*F*(2,22) = 51.38, *p* < .001, *p-Holm* < .001], but neither a main effect of location nor attention repetition [*p* > .1], nor any two-way [*ps* > .1] or three-way [*p* > .1, *BF_10_* = 0.09] interactions.

Lastly, for the correctness analysis (**Figure 4D**), we conducted a 3-way rmANOVA on the factors of attention (valid, neutral, invalid), location (left, right, upper, lower), and *n* – 1 correctness (correct, incorrect). Results revealed a main effect of attention [*F*(2,22) = 48.93, *p* < .001, *p-Holm* < .001], but neither a main effect of location nor *n –*1 correctness [*p* > .1], nor any two-way [*ps* > .1] or three-way [*p* > .1, *BF_10_* = 0.23] interactions.

In addition, we analyzed the data separately by the first and second halves of the blocks within each session and between the first and second sessions, we observed no interactions among attention, location, and any sequential effect (*ps* > .1), supporting the lack of interaction between trial history and attentional effects across locations. To summarize, an effect of attention was observed across all analyses: performance was higher in the valid condition and lower in the invalid condition than in the neutral condition. However, there were no significant effects of location or sequential history, nor any interactions involving target location.

#### Study 2

For the location analysis (**Figure 5A**), results revealed a main effect of attention [*F*(2,16) = 72.69, *p* < .001, *p-Holm* < .001], but neither a main effect of location nor location repetition [*ps* > .1], nor any two-way [*ps* > .05] or three-way [*p* > .1, *BF_10_* = 0.06] interactions.

**Figure 5.**
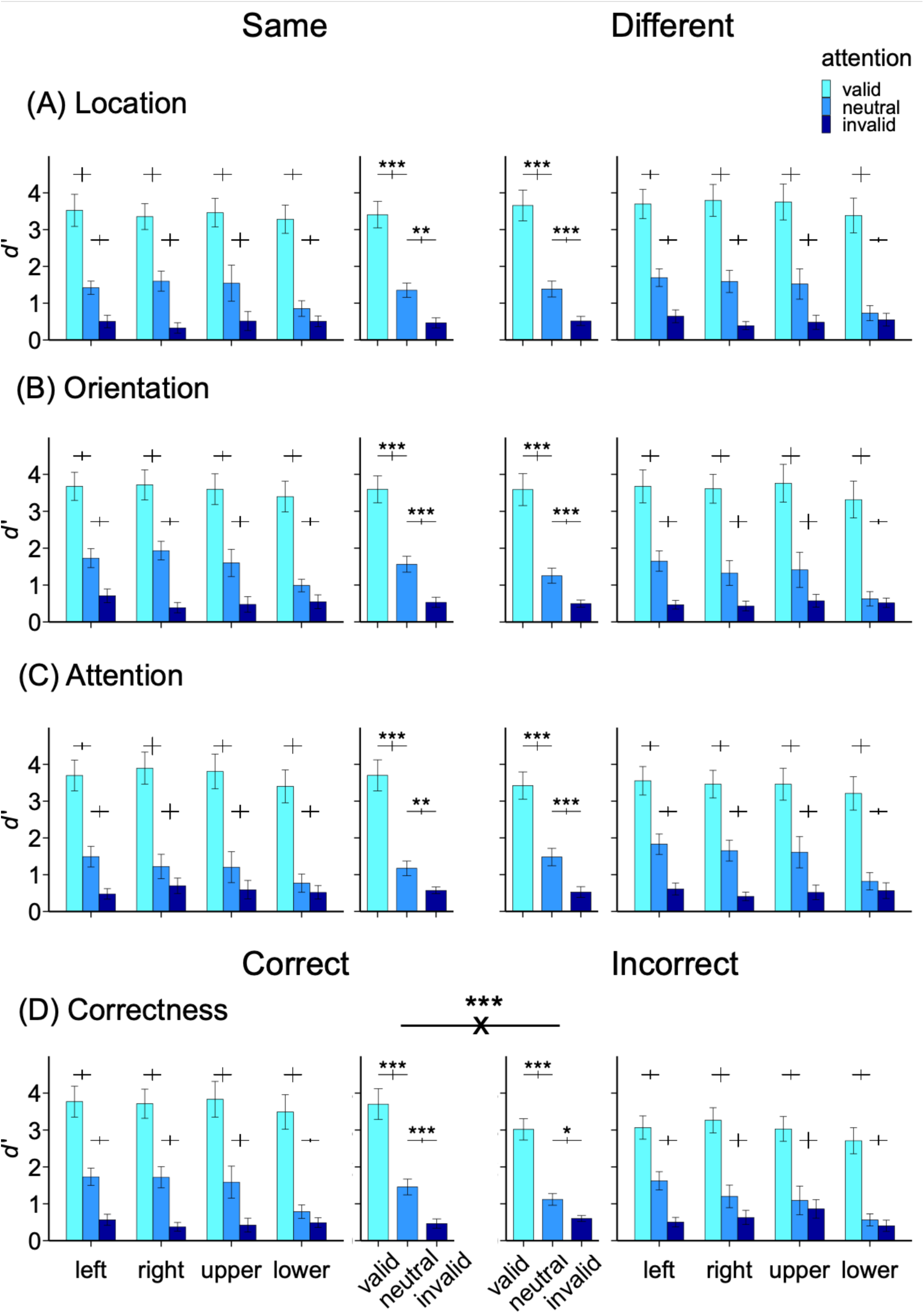
Study 2. Sensitivity in the orientation discrimination task when the (A) location, (B) target’s orientation, or (C) attentional cueing condition in the previous (*n –* 1) trial was the same as or different from the current trial. (D) Sensitivity when the performance on the previous (*n –* 1) trial was correct or incorrect. For each panel, left and right bars represent the same and different (or correct and incorrect) conditions, respectively. The middle two columns indicate the results averaged across locations. Error bars within the bars depict ±1 *SEM* of the condition; error bars above the bars indicate ±1 *SEM* of the difference between conditions.

For the feature analysis (**Figure 5B**), results revealed a main effect of attention [*F*(2,16) = 72.92, *p* < .001, *p-Holm* < .001], but no main effects of location or orientation repetition [*ps* > .1]. The two-way interaction between attention and orientation repetition was no longer significant after correction for multiple comparisons [*F*(2,16) = 4.44, *p* = .029, *p-Holm* = .059]. There were no other two-way [*ps* > .05] or three-way [*p* > .1, *BF_10_* = 0.06] interactions.

For the attention analysis (**Figure 5C**), results revealed a main effect of attention [*F*(2,16) = 72.11, *p* < .001, *p-Holm* < .001], but no main effect of location or attention repetition [*ps* > .1].

There was a two-way interaction between attention and attention repetition but not after corrections for multiple comparisons [*F*(2,16) = 5.32, *p* = .017, *p-Holm* = .051]. There were no other two-way [*ps* > .1] or three-way [*p* > .1, *BF_10_* = 0.07] interactions.

For the correctness analysis (**Figure 5D**), results revealed a main effect of attention [*F*(2,16) = 77.93, *p* < .001, *p-Holm* < .001], but no main effect of *n –*1 correctness after multiple-comparison correction [*F*(1,8) = 7.34, *p* = .027, *p-Holm* = .106], nor of location [*p* > .05]. There was a two-way interaction between attention and *n* –1 correctness [*F*(2,16) = 11.51, *p* < .001, *p-Holm =* .003], in which *d′* in the valid condition was higher following a correct trial than an incorrect trial [*t*(8) = 4.43, *p* = .002], but no differences were found in the neutral or invalid conditions [*ps* > .05]. There were no other two-way [*ps* > .1] or three-way [*p* > .1, *BF_10_* = 0.11] interactions.

To summarize, an effect of attention was observed across all analyses: performance was higher in the valid condition and lower in the invalid condition than in the neutral condition. Moreover, sensitivity increased after correct, compared to incorrect trials, in the valid condition. No significant interactions were observed between location and sequential effects.

#### Study 3

For the location analysis (**Figure 6A**), results revealed main effects of attention [*F*(1,17) = 58.14, *p* < .001, *p-Holm* < .001] and location repetition [*F*(1,17) = 9.42, *p* = .007, *p-Holm* = .021], in which non-repeated trials had a higher *d′* than repeated ones. There was neither a main effect of location [*p* > .1] nor any two-way [*ps* > .05] or three-way [*p* > .1, *BF_10_* = 0.12] interactions.

**Figure 6.**
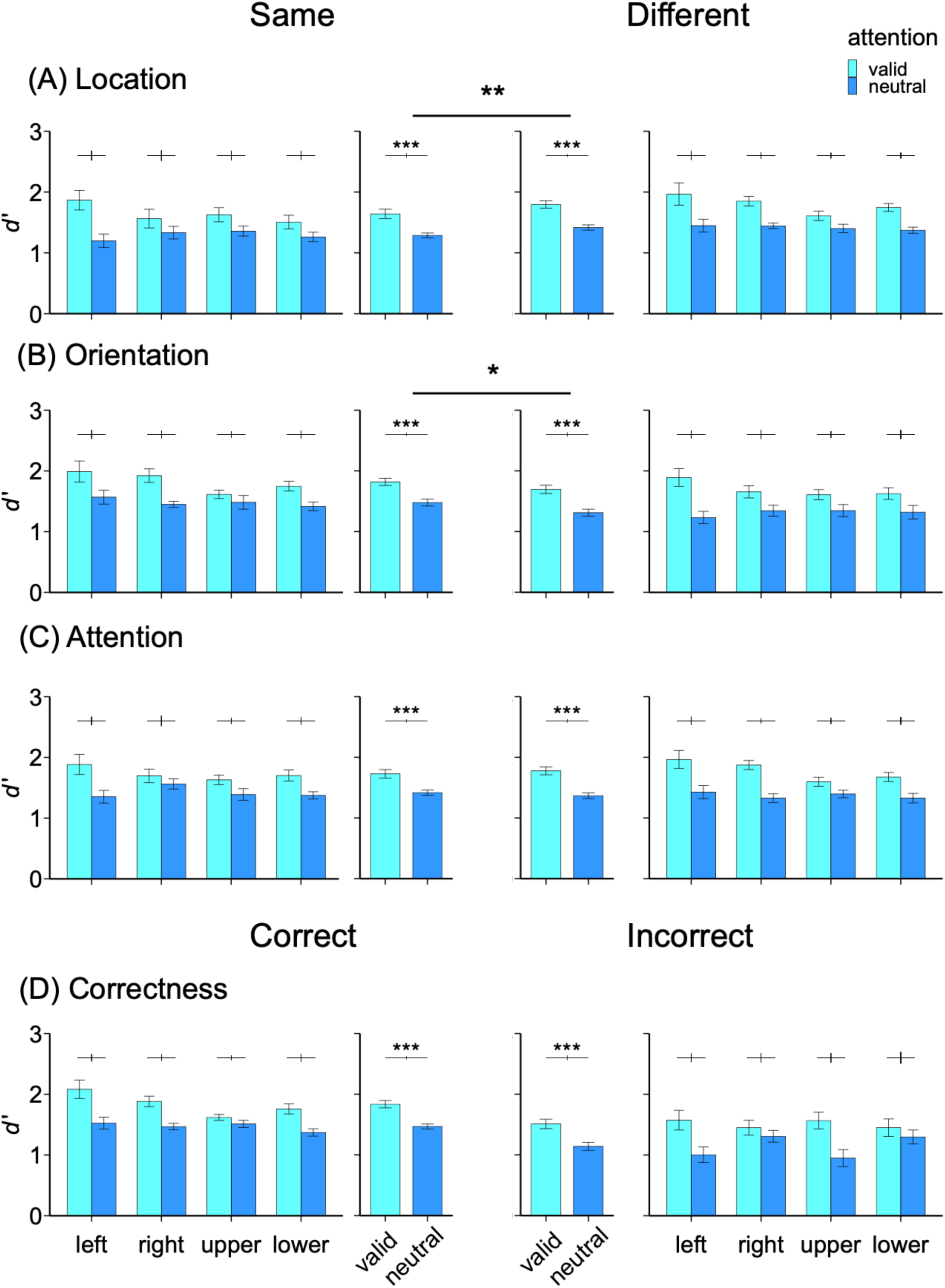
Study 3. Sensitivity in the orientation discrimination task when the (A) location, (B) target’s orientation, or (C) attentional cueing condition in the previous (*n –* 1) trial was the same as or different from the current trial. (D) Sensitivity when the performance on the previous (*n –*1) trial was correct or incorrect. For each panel, left and right bars represent the same and different (or correct and incorrect) conditions, respectively. The figures in the middle two columns indicate the results averaged across locations. Error bars within the bars depict ±1 *SEM* of the condition; error bars above the bars indicate ±1 *SEM* of the difference between conditions.

For the feature analysis (**Figure 6B**), results revealed main effects of attention [*F*(1,17) = 79.43, *p* < .001, *p-Holm* < .001] and orientation repetition [*F*(1,17) = 6.39, *p* = .022, *p-Holm* = .043], in which repeated trials had a higher *d′* than non-repeated ones. There was neither a main effect of location [*p* > .1] nor any two-way [*ps* > .05] or three-way [*p* > .1, *BF_10_* = 0.17] interactions.

For the attention analysis (**Figure 6C**), results revealed a main effect of attention [*F*(1,17) = 81.45, *p* < .001, *p-Holm* < .001], but neither a main effect of location nor attention repetition [*ps* > .1], nor any two-way [*ps* > .1] or three-way [*p* > .1, *BF_10_* = 0.33] interactions.

For the correctness analysis (**Figure 6D**), results revealed main effects of attention [*F*(1,17) = 59.65, *p* < .001, *p-Holm* < .001] and of *n –*1 correctness [*F*(1,17) = 29.28, *p* < .001, *p-Holm* < .001], but not of location [*p* > .1]. There was a three-way interaction [*F*(3,51) = 3.92, *p* = .014, *p-Holm* = .041]. Paired *t*-tests showed that in the *n –* 1 correct condition, the attentional benefit at the upper vertical meridian was smaller than at both the left [*t*(17) = 3.37, *p* = .004] and right [*t*(17) = 2.19, *p* = .043] horizontal meridians, but no other comparisons were significant [*ps* > .1]. In the *n –* 1 incorrect condition, in contrast, the attentional benefit at the upper vertical meridian was greater than at both the right horizontal [*t*(17) = 2.56, *p* = .02] and lower vertical meridians [*t*(17) = 2.27, *p* = .037], but no other comparisons were significant [*ps* > .05].

In sum, there was an effect of attention across all analyses, whereby performance was higher in the valid condition, followed by the neutral and invalid conditions. In general, sensitivity (*d′*) was higher when the previous target appeared at a different location and when the previous trial shared the same orientation. Whereas no interactions between location and sequential effects emerged in the location, feature, or attention analyses, a significant interaction was found in the correctness analysis. Specifically, performance at the upper vertical meridian was lower than at other locations when the previous trial was correct, but higher when the previous trial was incorrect.

#### Combined analysis for performance

### The attention benefit

To examine the consistency of our findings across studies and enhance statistical power, we combined the data from the three studies and conducted four analyses. We treated the attentional benefit (valid *d′* – neutral *d′*) as the dependent variable and conducted a series of 2-way rmANOVAs on the factors of location (left, right, upper, lower), and trial history’s (*n* –1) location, orientation, attention (same, different), or correctness (correct, incorrect).

For the location (**Figure 7A**, top panel), feature (**Figure 7B**, top panel), attention repetition (**Figure 7C**, top panel) and *n* – 1 correctness analyses (**Figure 7D**, top panel), results revealed no main effects of location [*ps* > .05] or trial history factors [*ps* > .1], nor any two-way interactions [*ps* > .1, all *BF_10_* < 0.33]. These results further show that the attentional benefit was stable and did not vary around the visual field, nor did it vary with the history of the prior trial.

**Figure 7.**
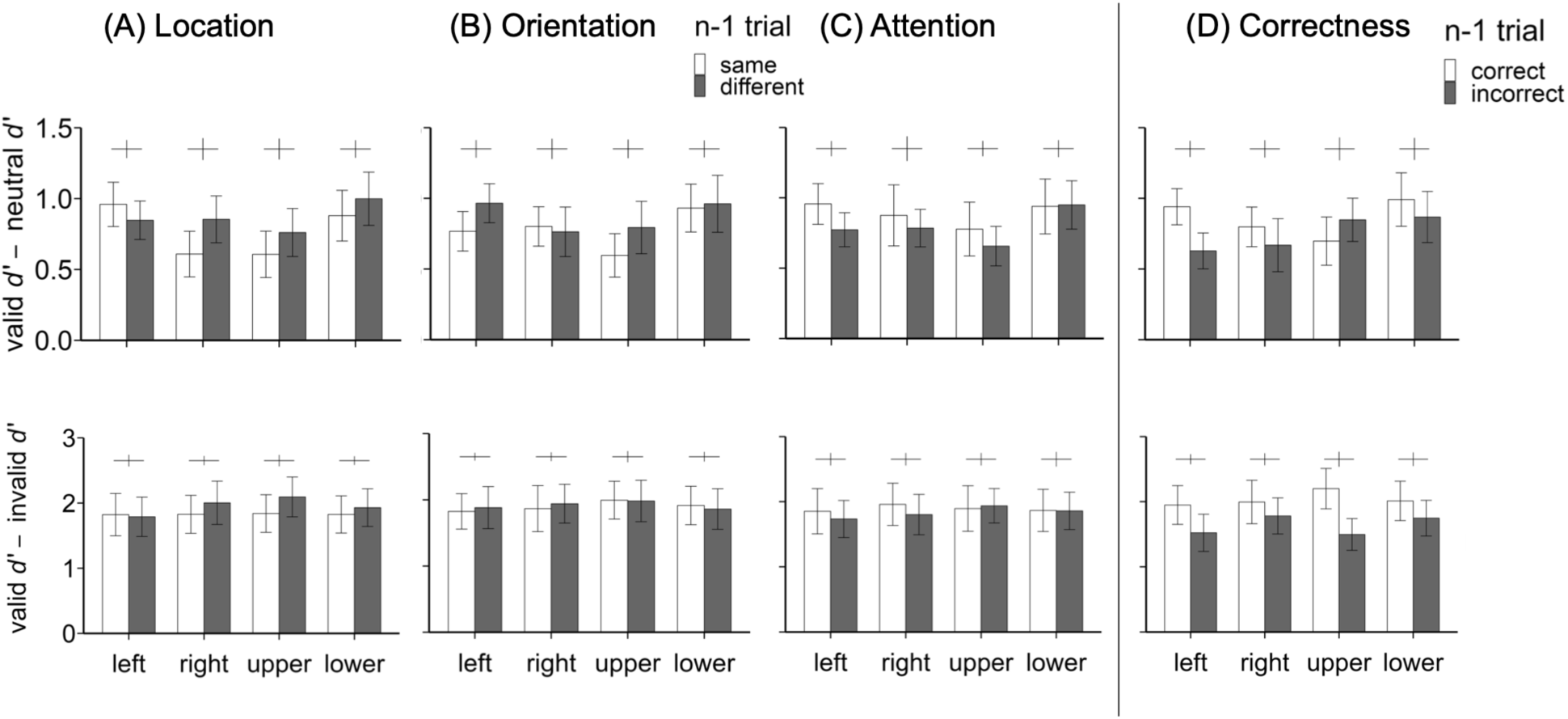
Combined analysis across Studies 1–3. The attentional benefit (top panels) and overall attention effect (bottom panels) in the orientation discrimination task when the (A) location, (B) target’s orientation, or (C) attentional cueing condition in the previous (*n –* 1) trial was the same as or different from the current trial. (D) The attentional benefit and overall attention effect when the performance on the previous (*n –*1) trial was correct or incorrect. The top panels depict the attentional benefit and the bottom panels depict the overall attentional effect. Error bars within the bars depict ±1 *SEM* of the condition; error bars above the bars indicate ±1 *SEM* of the difference between conditions.

### The overall attention effect

To examine the consistency of attentional modulation, as reflected by the overall attention effect, we evaluated the validity effect (valid *d′* - invalid *d′*) across Studies 1 and 2 that contained invalid trials. We conducted a series of 2-way rmANOVAs on the factors of target location (left, right, upper, lower), and trial history’s (*n* –1) location, orientation, attention (same, different), or correctness (correct, incorrect).

For the location (**Figure 7A**, bottom panel), feature (**Figure 7B**, bottom panel), and attention repetition analyses (**Figure 7C**, bottom panel), results revealed no main effects of repetition or location [*ps* > .05] nor any two-way interactions [*ps* > .1, all *BF_10_* < 0.1]. These results further demonstrate that the overall attention effect was stable, regardless of the location, orientation, or attentional condition in the previous trial.

For the correctness analysis (**Figure 7D**, bottom panel), there was a main effect of *n* – 1 correctness, in which the overall attentional effect was higher when the previous trial was correct than when it was incorrect [*F*(1,20) = 12.54, *p* = .002, *p-Holm* = .004], but neither a main effect of location [*p* > .1] nor an interaction emerged [*p* > .1]. Thus, previous correct trials led to an overall larger attention effect than previous incorrect trials.

### The standard deviation of reaction time

In addition to sensitivity and attentional effects, we further examined whether sequential effects modulate response variance, as previous studies found that the error distribution becomes smaller under sequential effects (Cicchini et al., 2018; Ozkirli et al., 2026; Sheehan & Serences, 2022). Given that our tasks were not reproduction tasks, we analyzed the standard deviation of reaction time (RT SD) to infer response variance. We excluded incorrect trials and reaction times exceeding ± 3*SD* from the mean of each participant. When pooling all three studies, we observed no main effects (*ps* > .1) nor any interactions among trial history (location, orientation, attention repetition, and *n* – 1 correctness), attention, and location (all *ps*>.05).

## Discussion

In all three studies analyzed here (Lee & Carrasco, 2026; Purokayastha et al., 2021; Tünçok, Carrasco, & Winawer, 2025), we assessed contrast response functions and contrast sensitivity using an orientation discrimination task, which is contingent upon stimulus contrast (Nachmias, 1967); a population-coding model has linked the neural response to psychophysical performance in orientation discrimination (Pestilli et al., 2009). Performance was quantified using signal detection theory, where *d’* indicates sensitivity to the orientation. Sensitivity (*d’*) is proportional to the signal-to-noise ratio of the underlying neuronal responses in the visual cortex. We observed a sequential effect in participants’ tendency to give a repeated response when the previous trial (*n –* 1) shared the same location or orientation as the current trial (**Figure 2**), consistent with previous findings (e.g., Can & Collins, 2025; Murai & Whitney, 2021; St. John-Saaltink et al., 2016). Critically, endogenous attention significantly modulated performance to a similar degree around polar angle, and sequential effects did not influence the relation between attention and location. Thus, although the sequential effect biased the response toward the repeated target location and feature (**Figures 2 and 3**), it did not improve performance across locations in endogenous attention tasks (**Figures 4–7**).

### Typical Sequential Effects in Response Repetition

For the response repetition analyses, participants were more likely to repeat their response if the previous target was at the same location as the current one (**Figures 2A and 3A**). This aligns with the spatial dimension of sequential effects, in which stronger biases occur when consecutive targets overlap in space (Bliss et al., 2023; Collins, 2019; Fischer & Whitney, 2014; Fornaciai & Park, 2018; Kondo et al., 2022; Luo et al., 2022; St. John-Saaltink et al., 2016) Additionally, a robust interaction emerged when the previous trial shared the same feature as the current one across three studies (**Figures 2B and 3B**; Barbosa & Compte, 2020; Fischer & Whitney, 2014; Liberman et al., 2014; Turbett et al., 2021). Specifically, when the previous trial had the same orientation and the current trial was in a valid condition, participants were more likely to repeat their response than when orientations differed. Interestingly, this relation was modulated by attentional condition; for the first time, we observed that the response repetition probability difference between the same and different orientations declined as a function of cue validity (**Figure 3B**). Additionally, the permutation test in the combined data analysis of valid and neutral trials confirmed that the probability difference in repeating responses on the same than different orientation trials was not due to the target feature itself but reflected sequential effects. These findings suggest that participants adaptively tailored their strategy in response to task demands.

When the attentional condition was repeated from the previous trial, it did not modulate the probability of giving a repeated response, except for the invalid condition when analyzing Studies 1 and 2 together (**Figure 3C**). Note that our analysis evaluated whether the attentional cueing condition was repeated, rather than its validity *per se*. Thus, participants did not change their response strategy in valid and neutral conditions but did so in the invalid condition: if the current trial was invalid, participants were more likely to rely on their previous response if the previous trial was from a different attentional condition.

As for trial correctness, more repeated responses were given if the previous trial was correct than incorrect (**Figure 3D**), a strategy participants adopted regardless of attentional condition. A correct response usually reinforces response repetition (Little & Clifford, 2025; St. John-Saaltink et al., 2016). However, this reinforcement did not benefit performance. Indeed, sequential effects on performance (sensitivity) were largely absent across all three studies, despite minor differences. Although errors can sometimes elicit compensatory adjustments that improve subsequent performance, post-error improvement in accuracy does not always occur (Adkins et al., 2024; Danielmeier & Ullsperger, 2011; Hajcak et al., 2003; Hajcak & Simons, 2008; Ullsperger et al., 2014): because post-error adjustments in accuracy depends on the number of trials and errors in an experiment and exhibit substantial individual differences, they are not always observed following an erroneous trial (Danielmeier & Ullsperger, 2011).

### Lack of Sequential Effects in Sensitivity

For the performance analyses, in Study 1, neither location nor its sequential history influenced performance. In Study 2, neither location nor target orientation history affected performance. In Study 3, sensitivity increased when the previous target appeared at a different location or when the previous trial shared the same orientation; however, neither effect interacted with attention or location. Notably, performance at the upper vertical meridian was lower following correct trials but higher following incorrect trials. Additionally, a combined analysis across the three studies confirmed that the attention effects were invariant across polar angles and did not interact with sequential effects.

These results, supported by Bayes Factor analyses, provide the first evidence that endogenous covert spatial attention is invariant not only to location but also to sequential effects. Although endogenous attention is flexible (for reviews, Carrasco, 2011; Carrasco & Barbot, 2014), here, it did not allocate more resources to locations where discriminability is intrinsically poor (i.e., the vertical meridian, especially the upper vertical meridian), even when considering previous trial conditions and response correctness. Thus, the finding that endogenous attention enhances contrast sensitivity similarly around polar angle is robust across eccentricities (from 6° to 8°), target tilt angles (from 2.5° to 20° tilt), target orientations (horizontal or vertical), spatial frequencies (from 4 to 5 cpd), sizes (from 3° to 4° of visual angle), and number of possible target locations (from 2 to 4) in these three studies (Lee & Carrasco, 2026; Purokayastha et al., 2021; Tünçok, Carrasco, & Winawer, 2025).

The finding that polar angle asymmetries are not modified by sequential effects aligns with their resilience to several fundamental visual processes. These include endogenous (Lee & Carrasco, 2026; Purokayastha et al., 2021; Tünçok, Carrasco, & Winawer, 2025) and exogenous (Cameron et al., 2002; Roberts et al., 2016, 2018) spatial attention, endogenous temporal attention (Fernández et al., 2019) and presaccadic attention, the latter of which even exacerbates these asymmetries (Hanning et al., 2022, 2024; Kwak et al., 2025; Kwak, Zhao, et al., 2024). However, recently, two visual phenomena—preview and adaptation—have been shown to alleviate performance asymmetries and render perception more homogeneous. First, the preview benefit is inversely related to polar angle asymmetries; it is largest at the upper vertical meridian and smallest at the horizontal meridian, indicating that the preview benefit helps compensate for performance asymmetries by integrating information across saccades (Liu et al., 2024). Second, the adaptation effect is stronger along the horizontal meridian than the vertical meridian, mitigating the horizontal-vertical asymmetry (HVA) in contrast sensitivity (Lee & Carrasco, 2025).

The present lack of sequential effects on performance in our orientation discrimination tasks may appear inconsistent with the stronger sequential effect in response bias observed in the periphery than in foveal vision in an adjustment task (Kandemir & Olivers, 2026). These authors propose that sequential effects scale inversely with perceptual sensitivity due to spatially-tuned continuity fields and spatial pooling in the visual cortex (Fischer & Whitney, 2014; Manassi et al., 2023; Manassi & Whitney, 2024). Accordingly, we would have expected stronger sequential effects at the upper vertical meridian where sensitivity is lowest. Our data, however, show no such amplification in performance, supporting the idea that eccentricity-dependent findings do not generalize to polar angle asymmetries in performance, which appear to be governed by distinct system-level computations (Xue et al., 2024; Xue, Barbot, et al., 2026; Xue, Landy, & Carrasco, 2026). Thus, this apparent discrepancy underscores the dissociation between sequential response bias (which favors repeated target locations and features) and perceptual performance (which remains invariant to sequential history).

Combining data from three studies examining endogenous attentional effects across the visual field, we gained higher statistical power and identified consistent patterns—such as a general absence of overall sequential effects on performance—despite parameter variations, such as the number of target locations, orientations, stimulus size, spatial frequency, eccentricity, inter-stimulus interval (ISI), and the inclusion of invalid trials. However, we also observed minor differences in the statistical outcomes across the individual protocols. In Study 2, higher performance occurred on valid trials following a correct trial than an incorrect one. When combining Studies 1 and 2, a correct preceding trial led to a larger overall attentional effect in the subsequent trial, regardless of target location (**Figure 7D**, bottom panel), a phenomenon likely driven by higher sensitivity on valid trials (**Figures 4D and 5D**), even though this effect did not reach statistical significance in Study 1. Thus, successful responses may reinforce the effective deployment of endogenous attention, enhancing its effect in the subsequent trial (Danielmeier & Ullsperger, 2011). In Study 3, we observed a sequential effect at the upper vertical meridian when analyzing performance following correct versus incorrect trials: a smaller attention benefit when the previous trial was correct, but a larger attention benefit when the previous trial was incorrect (**Figure 6D**). The protocol in Study 3 differed from Studies 1 and 2 in that cue validity was 100% (rather than 75%). Accordingly, participants were more likely to rely on the cue (Giordano et al., 2009; Kinchla, 1980; Mangun & Hillyard, 1990; Sperling & Melchner, 1978; Vossel et al., 2006). We speculate that participants reweighted their attention resources nonuniformly following an error, with the upper vertical meridian receiving more resources. Future research could systematically manipulate these varying parameters to precisely determine the specific factors responsible for these minor differences.

Overall, we observed sequential effects on response repetition probability but did not observe sequential effects on performance regarding location, target orientation, attention repetition, or *n* – 1 correctness. When examining sequential effects or serial dependence, most studies have reported how the response was biased toward the previous stimulus, via adjustment or reproduction tasks (e.g., Ceylan & Pascucci, 2023; Kondo et al., 2022; Manassi et al., 2023; Samaha et al., 2019). Whereas we confirm this response repetition bias using orientation discrimination tasks (e.g., Can & Collins, 2025; Murai & Whitney, 2021; St. John-Saaltink et al., 2016), perceptual ‘sensitivity’ or ‘performance’ has rarely been evaluated alongside it. Thus, sequential effects on performance and on response repetition should not be directly compared. The current study revealed that neither the magnitude of endogenous spatial attentional benefits nor overall sensitivity was modulated by prior trial history across spatial, feature, or attentional dimensions. These results are consistent with findings from an endogenous temporal attention orientation discrimination task: perceptual sensitivity was independent of the temporal attentional condition in the previous trial (Duyar & Carrasco, 2025a).

Using a two-alternative forced choice (2AFC) task, rather than a reproduction task, to evaluate sequential effects presents certain limitations. First, we could not evaluate the attraction versus repulsion effect, which has been widely investigated in studies of sequential effects, in which the direction and underlying mechanisms remain an open question (e.g., Fritsche et al., 2017; Gallagher & Benton, 2022; Pascucci et al., 2019). Second, because responses were categorical rather than continuous, we could not assess the magnitude or distribution of response errors across trials, as in reproduction tasks (e.g., Cicchini et al., 2018; Ozkirli et al., 2026; Sheehan & Serences, 2022). This distinction is relevant when considering whether sequential dependence provides a functional benefit to perceptual performance. In reproduction tasks, performance can be characterized not only by the mean response error or bias but also by the variability of response errors, providing an index of response precision, although whether serial dependence generally improves performance efficiency remains to be determined (Ozkirli et al., 2026), and our RT *SD* analysis did not show effects of trial history. By contrast, 2AFC tasks provide complementary advantages. In an orientation discrimination task, performance can be systematically contingent on stimulus contrast for quantifying contrast sensitivity (Nachmias, 1967; Pestilli et al., 2009). Moreover, because participants make a categorical perceptual judgment rather than reproduce an exact stimulus value, 2AFC tasks may reduce sources of variability associated with continuous response adjustment and motor reproduction. Thus, whereas reproduction tasks provide fine-grained information about the direction and magnitude of trial-by-trial biases, the 2AFC task here allows us to assess whether previous-trial information systematically influences subsequent perceptual decisions and sensitivity.

In contrast to the lack of sequential effects on performance in our covert endogenous attention tasks, presaccadic attention exhibits sequential effects based on location and feature analysis in a motion discrimination task (White et al., 2013), and eye movement patterns are modulated by the previous trial in an orientation reproduction task (Tyralla & Zimmermann, 2025). Presaccadic attention differs from endogenous attention in its task structure (whether eye movements are executed), how each shapes sensory tuning (Li et al., 2016; Li et al., 2021; Li et al., 2019), and their neural underpinnings (Fernández et al., 2023; Hanning et al., 2023). Our findings provide further evidence that these two types of spatial attention yield different perceptual outcomes and sequential effects.

The neural underpinnings of sequential effects remain a subject of debate (Manassi & Whitney, 2024; Pascucci et al., 2023). Relevant to the current study are fMRI findings showing that in orientation discrimination tasks, signals in V1 are biased toward the orientation presented on the preceding trial (St. John-Saaltink et al., 2016; Wang et al., 2026), and TMS findings demonstrating that stimulating the lateral prefrontal cortex can effectively reduce serial dependence (Bliss et al., 2023). It is possible that early visual cortex receives top-down signals from the prefrontal area and contributes to serial dependence (Cicchini et al., 2021; Manassi & Whitney, 2024; Wang et al., 2026). For endogenous attention, another top-down mechanism modulates fMRI BOLD signals in the early visual cortex: higher-order, frontoparietal attentional regions send feedback information to the visual cortex, with effects diminishing in earlier visual areas (Beck & Kastner, 2014; Chica et al., 2013; Dugué et al., 2020; Lauritzen et al., 2009; Pestilli et al., 2011; Puckett & DeYoe, 2015; Tünçok, Carrasco, & Winawer, 2025). Moreover, TMS studies have revealed that the human homolog of the right frontal eye fields (rFEF+) is functionally necessary for endogenous attention, but early visual cortex is not (Fernández et al., 2023).

In contrast, polar angle asymmetries have been linked to optical and retinal factors (Kupers et al., 2019, 2022) and to the functional architecture of the early visual cortex (Benson et al., 2021; Himmelberg et al., 2021, 2022, 2025). Although cortical surface area contributes substantially and correlates with contrast sensitivity and acuity (Benson et al., 2021; Himmelberg et al., 2022), it does not fully account for these asymmetries (Himmelberg et al., 2023; Jigo et al., 2023). Additional factors, such as variations in the gain of sensory tuning, also play a role (Xue, Barbot, et al., 2026; Xue, Landy, & Carrasco, 2026).

Whether serial dependence operates at the perceptual level (Burr & Cicchini, 2014; Cicchini et al., 2017; Collins, 2020; St. John-Saaltink et al., 2016) or at the post-perceptual decision level (Alais et al., 2017; Bliss et al., 2017; Ceylan et al., 2021; Fritsche et al., 2017) remains debated. Although not our primary focus, our findings support a decision-level account given that trial history altered response repetition probability but performance (*d′*) remained invariant. This creates a dissociation between perception and response strategy during endogenous attention tasks.

In conclusion, by analyzing three datasets of endogenous attentional effects across four different aspects of trial history, we demonstrate typical sequential effects on response repetition bias. However, despite its flexible nature, endogenous attention does not preferentially enhance contrast sensitivity at any specific location, even when accounting for the location, feature, attention, and correctness of the preceding trial. These findings reveal that visual performance fields are not reshaped by covert spatial attention, regardless of recent trial history or top-down modulation, underscoring the robustness of performance asymmetries in human visual perception.

## Acknowledgments

This study was supported by NIH NEI R01-EY027401 to M.C. and the Ministry of Education in Taiwan and 2025 National Science and Technology Council (NSTC) Taiwanese Overseas Pioneers Grants (TOP Grants) for PhD Candidates to H.-H.L. We thank David Tu, Katia Steinfeld, Marc Himmelberg, and Shutian Xue for their helpful feedback.

