## Supplementary Materials for "Endogenous attention is invariant to sequential effects in performance around the visual field"

To examine whether different trial history factors interact with one another and with the attentional condition, we conducted a 4-way rmANOVA on location repetition, orientation repetition, attention repetition and attention condition (valid, neutral) by pooling data across all three studies. We observed no significant 4-way interaction [$F(1,38) < 1$], but 3-way interactions among (1) location repetition, orientation repetition, and attentional condition [$F(1,38) = 4.91$, $p = .033$], and (2) orientation repetition, attention repetition, and attentional condition [$F(1,38) = 35.57$, $p < .001$].

For the location repetition × orientation repetition × attention condition interaction (**Figure S1A**), we observed no interaction between location and orientation repetition on valid trials [$F(1,38)<1$] but an interaction emerged on neutral trials [$F(1,38) = 11.76$, $p = .001$]: the location repetition difference is more striking when the orientation was the same than different.

For the orientation repetition × attention repetition × attention condition interaction (**Figure S1B**), we observed 2-way interactions on both valid [$F(1,38) = 32.58$, $p < .001$] and neutral [$F(1,38) = 14.75$, $p < .001$] trials. The difference in repetition proportion between same- and different-orientation trial varied as a function of the previous attentional condition: on valid trials, the attention repetition effect was more pronounced when orientations were repeated; whereas on neutral trials, this pattern reversed: the attention repetition effect was smaller when orientations were repeated.

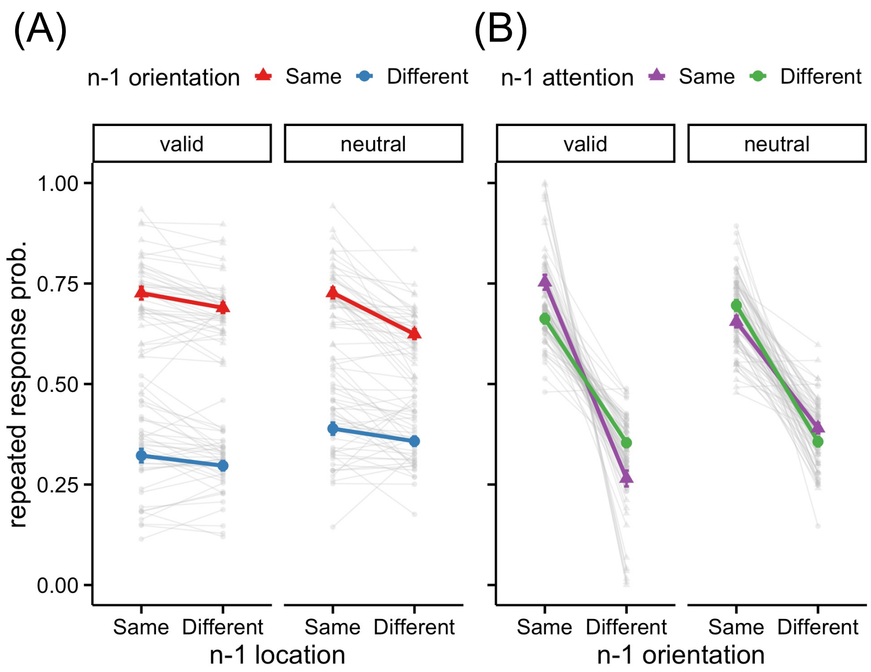


**Figure S1.** (A) Probability of giving a repeated response in the orientation discrimination task pooled across the three studies when (A) location and target's orientation were the same as or different from the previous (*n* – 1) trial in valid and neutral trials, or (B) target's orientation and attention were the same or different the previous (*n* –1) trial on valid and neutral trials. Error bars within the bar plots depict ±1 *SEM* of the condition. Gray lines indicate data from individual participants.
